# Cortical microinfarcts potentiate recurrent ischemic injury through NLRP3-dependent trained immunity

**DOI:** 10.1101/2020.06.17.157289

**Authors:** Yiwei Feng, Tengteng Wu, Yukun Feng, Fengyin Liang, Ge Li, Yongchao Li, Yalun Guan, Shuhua Liu, Yu Zhang, Guangqing Xu, Zhong Pei

**Affiliations:** Department of Neurology, The First Affiliated Hospital, Sun Yat-sen University, Guangzhou, 510080, China; Guangdong Provincial Key Laboratory of Diagnosis and Treatment of Major Neurological Diseases, National Key Clinical Department and Key Discipline of Neurology, Guangzhou, 510080, China; Guangdong Provincial Key Laboratory of Laboratory Animals, Guangdong Laboratory Animals Monitoring Institute, Guangzhou, 510080, China; Department of Rehabilitation Medicine, Beijing Tiantan Hospital, Capital Medical University, Beijing, 100070, China; China National Clinical Research Center for Neurological Diseases, Beijing, 100070, China; Department of Neurology, Huashan Hospital, Fudan University, Shanghai, 200040, China; Department of Neurology, Zhongshan Hospital, Fudan University, Shanghai 200032, China; Department of Neurology, Hainan General Hospital, 570311, Hainan, China

**Keywords:** NLRP3, microinfarct, recurrent stroke, Trained immunity

## Abstract

Microinfarcts are common among the elderly, and patients with microinfarcts are more vulnerable to another stroke. However, the potential effect of microinfarct on recurrent stroke remains elusive. In this study, we investigated the detrimental effect of microinfarct on recurrent stroke in mice. Microinfarct was induced using two-photon laser and photothrombotic stroke was induced in the cortex contralateral to microinfarct four weeks later. We found that CMI could trigger the formation of innate immune memory, which exacerbated the pro-inflammatory response and ischemic injury in second photothrombotic stroke. Furthermore, we clarified the role of NLRP3 inflammasome in the nuclei of microglia, which interacts with the MLL1 complex and thereby increases H3K4 methylation, suggesting that NLRP3 is critical in microinfarct-induced innate immune memory. Additionally, NLRP3 knockout in microglia attenuated microinfarct-induced detrimental effects on recurrent stroke. Our study highlights the detrimental effect of trained immunity on the recurrent stroke and reveals the important role of NLRP3 in mediating the formation of this memory, which may be a therapeutic target to mitigate recurrent strokes.

## Introduction

Strokes are the leading cause of death worldwide. Cortical microinfarcts (CMIs) are common and patient with microinfarcts are at high risk of recurrent strokes (15 to 42% over 5 years)(WilsonAmbler et al., 2019). Recurrent strokes account for up to 40% of all strokes and are associated with high mortality and poor functional recovery(Ovbiagele & Nguyen-Huynh, 2011). However, the influence of primary CMIs on the risk of a secondary stroke remains unclear.

Inflammation and immune mediators play a key role in stroke pathogenesis(Macrez, Ali et al., 2011). NLRP3 inflammasome, a multi-protein complex, is heavily involved in brain inflammation(Guo, Callaway et al., 2015, Mullard, 2019). Furthermore, high NLRP3 expression has been detected in the ischemic brain of stroke patients and animals, while NLRP3 inhibition alleviates ischemic brain damage(Hong, Gu et al., 2019, Ismael, Zhao et al., 2018). Our previous works has found that inhibition of NLRP3 significantly reduces recurrent stroke, suggesting that NLRP3 is an important molecule in recurrent stroke(He, Zeng et al., 2020).

The inflammatory response in the brain is mainly mediated by microglia, highly plastic, and phenotypically diverse cells which adapt to their microenvironment through different signal pathways (Delpech, Madore et al., 2015). Recently, microglia have been reported to play a key role in the development of trained immunity in the brain through histone modifications after inflammatory stimulation (Wendeln, Degenhardt et al., 2018). For example, enhanced H3K4me1/3 modification can open the chromatin and make the modified gene more easily to transcript(Netea, Joosten et al., 2016). Trained microglia produce excessive pro-inflammatory cytokines, which can damage brain cells in experimental models of neurological diseases (e.g., Alzheimer’s disease, stroke). Interestingly, it has been reported that NLRP3 also has a critical role in trained immunity(Christ, Günther et al., 2018).

Given that cerebral ischemia can induce epigenetic modifications in the brain, we investigated whether CMIs can induce trained immunity and a subsequent increase in the risk of recurrent strokes. We also examined the role of NLRP3 in the epigenetic reprogramming of H3K4me1 and H3K4me3, two key epigenetic marks of trained immunity, which provides an important therapeutic target to mitigate the consequences of recurrent stroke.

## Results

### 1.1 Effect of cortical microinfarct on innate immune memory in the contralateral brain cortex

Trained immunity in the brain is driven by the epigenetic changes in microglia. Given that cerebral ischemia can induce epigenetic alterations, we examined whether microinfarctions can promote epigenetic alterations(Qureshi & Mehler, 2011). A single CMI in left cortex was created by laser-induced occlusion of penetrating arteriole. Nissls staining revealed a highly localized small tissue infarction following CMIs in the left cortex, while CMIs did not cause any damage in contralateral cortex (Fig. 1a). We then examined the expression of H3K4me1 and H3K4me3, the active histone modifications, in microglia isolated from the cortex contralateral to the ischemic cortex following CMI. We found that the expression of H3K4me1 and H3K4me3 increased after the first week and remained high over the subsequent three weeks (Fig. 1b). H3K4me3 ChIP was performed to examine the role of H3K4me1 and H3K4me3 in the regulation of TNF-α, IL-6, and IL-1β. As expected, H3K4me1 and H3K4me3 modification was significantly increased on TNF-α, IL-6s and IL-1β promoter, indicating that microglia were epigenetically modified (Fig. 1c-e). However, TNF-α, IL-6, and IL-1β production remained unchanged over four weeks, indicating that histone modifications induced by CMI are not sufficient to alter the levels of TNF-α, IL-6, and IL-1β (Fig. 1f and g).

**Fig. 1.**
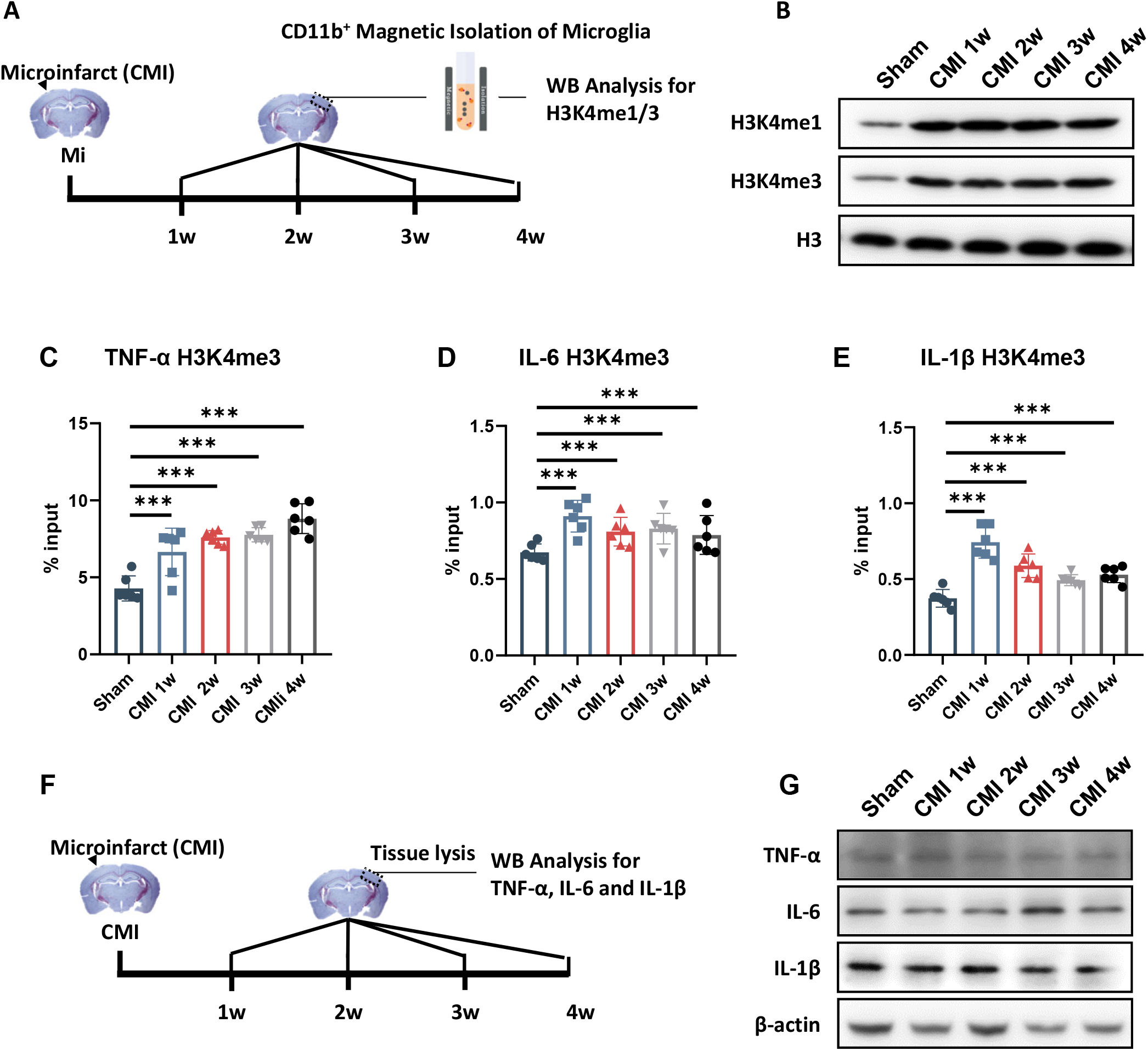
CMI-induced innate immune memory in the contralateral brain cortex. A Schematic diagram of the experimental design used in this study. B Immunoblots of H3K4me1/3 in the contralateral cortex after CMI induction. (N = 6 mice per group) C-E Quantitative analysis of H3K4me3 ChIP-qPCR on the TNF-α, IL-6, and IL-1β promoters in the contralateral microglia after CMI induction. (N = 6 mice per group) F Schematic diagram of the experimental design. G Immunoblots of TNF-α, IL-6, and IL-1β showing microglial pro-inflammatory cytokines expression in the contralateral cortex after CMI induction. (N = 6 mice per group) Results are presented as mean ± standard deviation (SD). All statistical tests were performed using two-way ANOVA and Sidak posthoc analysis. (ns, not significant; * *p* < 0.01; ** *p* < 0.01; *** *p* < 0.001).

### 1.2 Effect of H3 methylation on CMI-induced inflammatory response and recurrent stroke risk

To examine the impact of CMI-induced trained immunity on recurrent stroke, a photothrombotic stroke (PT) in the cortex contralateral to microinfarct was induced four weeks following initial microinfarction (Fig. 2a and b). Infarct size was significantly larger in mice with CMI than those without (Fig. 2c). Additionally, IL-6, IL-1β, and TNF-α production were significantly higher in the peri-infarct region in recurrent strokes (Fig 2d). IL-6, IL-1β, and TNF-α were also measured in collected blood samples to examine their possible effect on the peripheral inflammatory response. There was no significant difference in peripheral IL-6, IL-1β, and TNF-α between mice with recurrent and single photothrombotic stroke (Fig. S1h-j). This suggests that brain inflammation in recurring strokes is less likely to be mediated by peripheral inflammation.

**Fig. 2.**
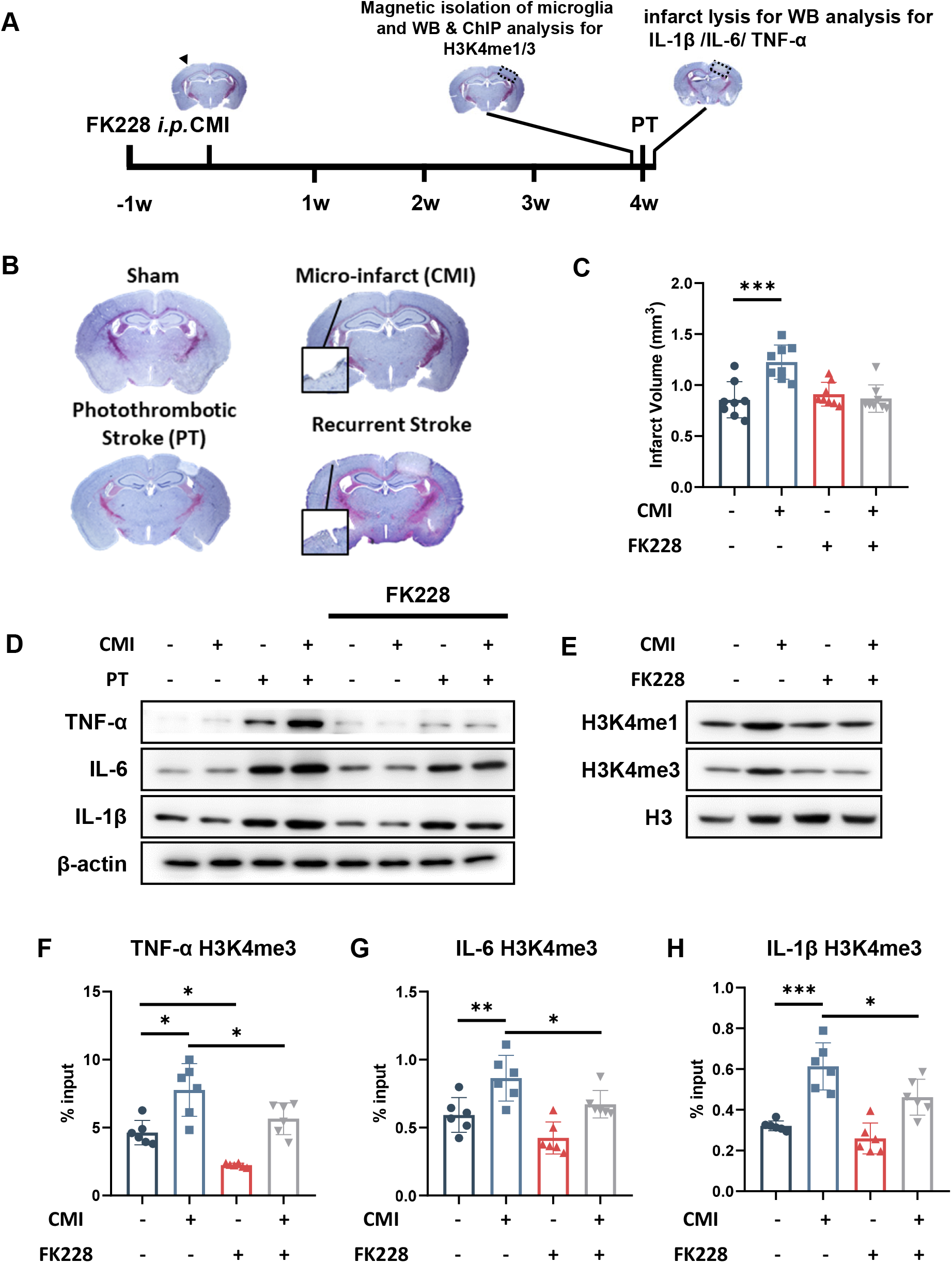
Effect of H3 methylation on CMI-induced inflammatory response and recurrent stroke risk. A Schematic diagram of experimental design. B, C Nissl staining (B) and quantitative analysis (C) showed the infarct size changes in photothrombotic stroke mice subjected to CMI and FK228 treatment (N = 6 mice per group) D Immunoblots of TNF-α, IL-6, and IL-1β showing pro-inflammatory cytokines expression in the perinfarct region 12 hours after the photothrombotic stroke (N = 6 mice per group). E Immunoblots of H3K4me1/3 showing microglial H3K4me1/3 expression in the contralateral cortex in mice subjected to CMI and FK228 treatment (N = 6 mice per group) F-H Quantitative analysis of H3K4me3 ChIP-qPCR on the TNF-α, IL-6, and IL-1β promoters in contralateral microglia after the induction of CMI and FK228 treatment (N = 6 mice per group) Results are presented as mean ± standard deviation (SD). All statistical tests were performed using two-way ANOVA and Sidak posthoc analysis. (ns, not significant; * *p* < 0.01; ** *p* < 0.01; *** *p* < 0.001).

To investigate the influence of histone modifications in the inflammatory response to CMI in a second photothrombotic stroke, HDAC inhibitor FK228, a major regulator of epigenetic reprogramming, was administrated a week before microinfarction. As expected, FK228 significantly inhibited expression of H3K4me1 and H3K4me3 (Fig. 2e) and the activation of H3K4me3 in TNF-α, IL-6, and IL-1β promoters (Fig. 2f-h). Similarly, FK228 attenuated the expression of TNF-α, IL-6, and IL-1β around the peri-infarct region after a thrombotic stroke (Fig. 2d).

Together, we demonstrated that CMI-induced trained immunity in mice, which in turn exacerbated ischemic damage in subsequent stroke.

### 1.3 Effect of CMI on the interaction of NLRP3 and MLL1 in the contralateral cortex

Recently, the NLRP3 inflammasome has been reported to mediate trained immunity(Christ et al., 2018). To investigate the role of NLRP3 inflammasome in trained immunity in recurrent stroke, we measured the expression of NLRP3 in microglia isolated from the cortex contralateral to the CMI. We found that NLRP3 expression robustly increased within the first two weeks and gradually returned to the baseline over the next two weeks (Fig. 3a). However, IL-1β production remained at the baseline level (Fig. 1g), indicating that the expression of NLRP3 did not promote IL-1β and IL-18 production. As NLRP3 is known to mediate the cleavage of IL-1β in the cytoplasm, we hypothesized that NLRP3 inflammasomes were primarily in microglial nuclei, and thus examined the localization of NLRP3 in isolated microglia. We found that NLRP3 expression was significantly higher in the nuclei within the first two weeks after CMI, whereas its expression in the cytoplasm remained unchanged (Fig. 3a). The nuclear localization of NLRP3 was further confirmed in immunofluorescent staining (Fig. 3b). To further investigate the link between NLRP3 and H3 methylation, we examined the interaction between NLRP3 and MLL1 (the main precursor of H3 methylation) by immunoprecipitation on microglia isolated from the non-ischemic cortex one week after CMI (Fig. 3c). There was no interaction between NLRP3 and MLL1 in the sham group but NLRP3 interacted with MLL1 in the microglia of CMI mice (Fig. 3c). The interaction of NLRP3 and MLL1 was further confirmed in microglial nuclei using a proximity ligation assay (Fig. 3d).

**Fig. 3.**
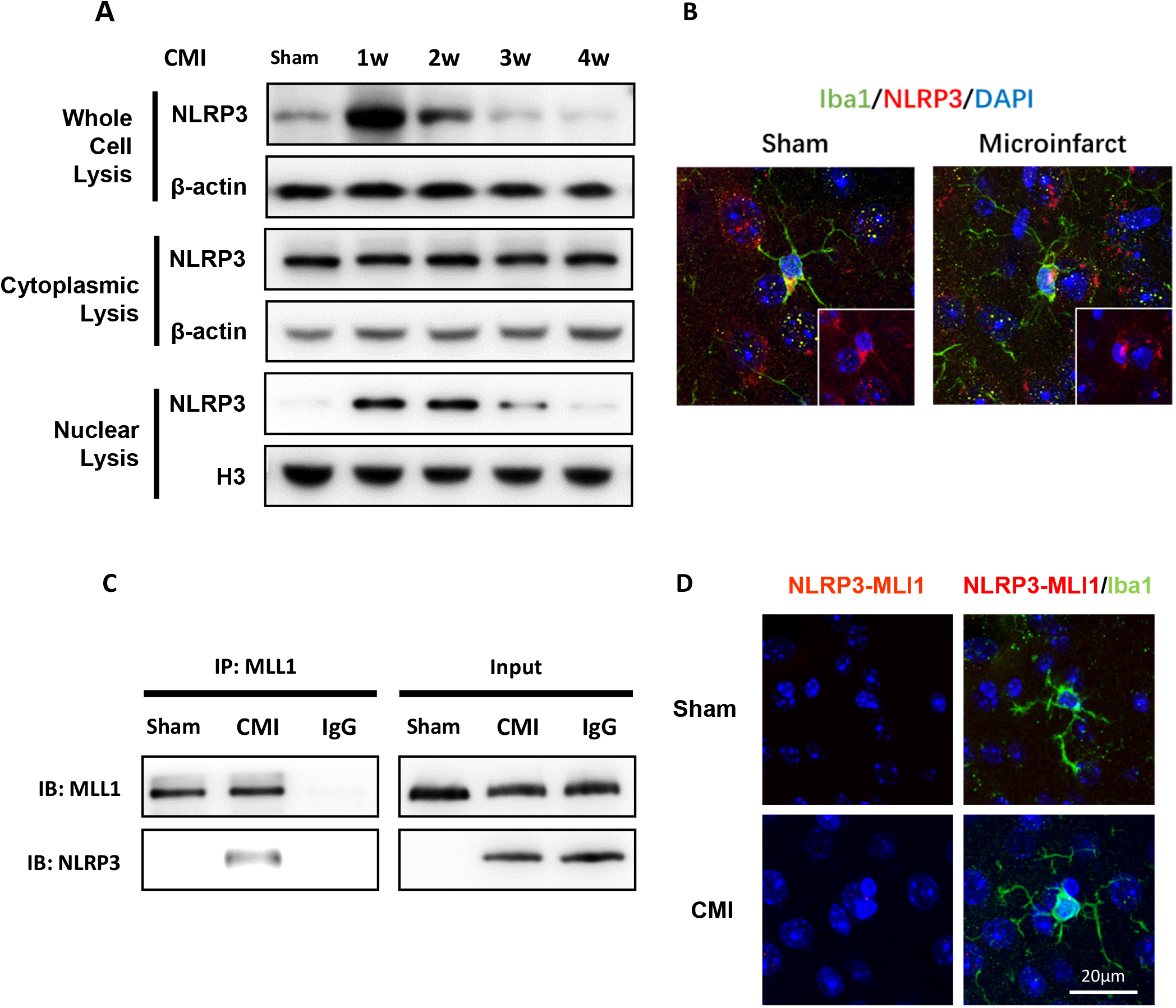
CMI-induced the interaction of NLRP3 with MLL1 in the contralateral cortex. A Immunoblots of NLRP3 expression in the microglial whole-cell lysis, cytoplasmic, and Nuclear lysis after CMI. (N = 6 mice per group) B Representative immunofluorescence of NLRP3 (red) expression in microglia (iba1, green) 1 weeks after CMI. C Immunoblot analysis of MLL1 or NLRP3 in the microglial lysates from mice subjected to sham and CMI, assessed after immunoprecipitation (IP) with anti-MLL1 or immunoglobulin control (Ig), or without immunoprecipitation (Input). D Proximity ligation assay of microglial (iba1, green) NLRP3 and MLL1 (red dots) in Sham and 1 week-post CMI mice Results are presented as mean ± standard deviation (SD). All statistical tests were performed using two-way ANOVA and Sidak posthoc analysis. (ns, not significant; * *p* < 0.01; ** *p* < 0.01; *** *p* < 0.001).

Collectively, NLRP3 interacted with MLL1 in microglial nuclei after CMI in the contralateral cortex, which in turn initiated the epigenetic re-programming of microglia and exacerbated the subsequent inflammatory response.

### 1. 4. Knockout of microglial NLRP3 mitigated MLL1, reduced H3 methylation, and attenuated CMI-induced detrimental effects on recurrent stroke

A microglial conditional knockout of NLRP3 (CX3CR1-CreER × NLRP3^fl/fl^ mice) was used to examine the functional influence of the interaction between the MLL1 complex and NLRP3 on recurrent stroke risk. The depletion of microglial NLRP3 was induced by tamoxifen administration four weeks before the microinfarction. MLL1 was significantly higher in WT microglial cells with CMI compared to those in the sham control (Fig. 4a) and we also observed high H3K4me1 and H3K4me3 expression in those cells (Fig. 4b), and an increase of H3K4me1 and H3K4me3 at TNF-α, IL-6 and IL-1β promoters (Fig. 4c-e). By contrast, CMI-induced alterations were reversed by the conditional knockout of NLRP3 (Fig. 4a-e). Most importantly, increased infarct size by CMI was mitigated in 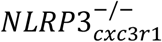 mice (Fig. 4f).

**Fig. 4.**
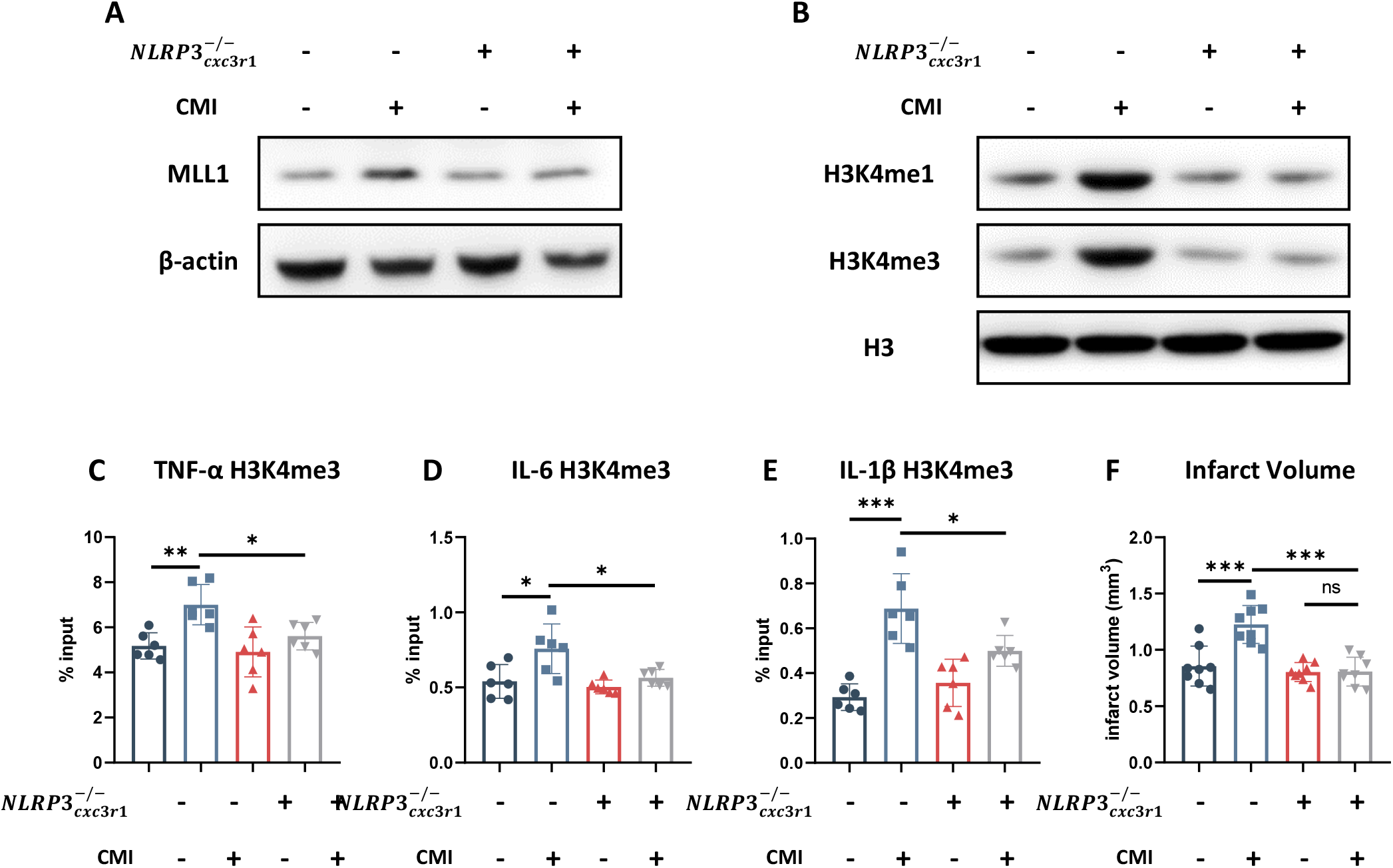
Knockout of microglial NLRP3 mitigated MLL1, reduced H3 methylation, and attenuated CMI-induced detrimental effects on recurrent stroke. A Immunoblot of MLL1 expression in WT and 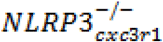 mice subjected to CMI treatment (N = 6 mice per group). B Immunoblot of H3K4me1/3 expression in WT and 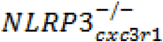 mice subjected to CMI treatment (N = 6 mice per group). C-E Quantitative analysis of H3K4me3 ChIP-qPCR on the TNF-α, IL-6, and IL-1β promoters among the contralateral microglia after the induction of CMI in WT and 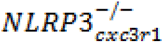 mice as indicated. (N = 6 mice per group) F Quantitative analysis of photothrombotic infarct volume in WT and 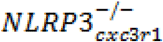 mice subjected to CMI (N = 6 mice per group) Results are presented as mean ± standard deviation (SD). All statistical tests were performed using two-way ANOVA with a Sidak posthoc analysis. (ns, not significant; * *p* < 0.01; ** *p* < 0.01; *** *p* < 0.001).

Taken together, our results show that NLRP3 in microglial nuclei was higher after CMI and subsequently interacted with the MLL1 complex, which promoted H3K4me1 and H3K4me3 expression. Therefore, NLRP3 may be a potential therapeutic target to mitigate the ischemic damage in recurrent strokes in CMI patients.

## Discussion

In this study, we found that CMI increases the risk of recurrent ischemic strokes through NLRP3-dependent trained immunity. The interaction of nuclear NLRP3 with MLL1 is critical in CMI-induced trained immunity, whereas knockout of microglia NLRP3 inhibits trained immunity and attenuates recurrent stroke. Our study highlights the importance of trained immunity in recurrent stroke risk and provides a novel insight of the influence of NLRP3 in this innate immunity memory formation.

CMIs are common among the elderly. However, recurrent stroke risk is difficult to predict, as most CMIs cannot be detected via magnetic resonance imaging. Recently, it has been reported that diffusion-weighted imaging can detect and monitor acute CMIs for a month, which may provide a better intervention window(Auriel, Westover et al., 2015, Ferro, van den Brink et al., 2019). In this study, we found that a single CMI could induce epigenetic alterations (H3K4me1 and H3K4me3) in the contralateral side, suggesting that a single CMI may have a big impact on the brain at the epigenetic level. This impact is significant, as it induces trained immunity, which leads to excessive immune activation and exacerbates the risk of subsequent strokes.

It is not clear how CMIs induce trained immunity. Traditionally, trained immunity is induced by vaccinations and inflammatory stimulators such as LPS(Mulder, Ochando et al., 2019). However, endogenous alarm signals associated with tissue damage and sterile inflammation can also induce trained immunity through the epigenetic regulation of histone modifications, such as damage-associated molecular patterns (DAMPs), oxLDL, and HDL(Crişan, Netea et al., 2016, Salam, Borsini et al., 2018). It has been reported that DAMPs activate TLR2 to epigenetically modify histone methylation and promote subsequent inflammatory responses. DAMPs and TLR2 are critical inflammatory signals involved in stroke-related inflammation, it is reasonable to speculate that trained immunity in the contralateral cortex of microinfarcts was induced similarly(Salam et al., 2018). Alternatively, released DAMPs from CMIs might reach the contralateral cortex through cerebrospinal fluid. The epigenetic modification caused by this damage signal further exacerbates the outcome of subsequent strokes.

Interestingly, transient cerebral ischemia (TIA) can induce protection against a subsequent stroke through preconditioning of brain cells (e.g., microglia), which differs from our observations in this study(Johnston, 2004, Sommer, 2009). TIA is a self-resolving focal cerebral ischemia with no evidence of acute brain tissue infarction(Johnston, 2004). As no substantial damage was observed in TIA, endogenous signals, such as DAMPs, may not be strong enough to initiate the inflammatory responses.

Histone modifications with chromatin reconfiguration are a central process in trained immunity(Netea et al., 2016). As a super-enhancer, H3K4me1/3 modifications unfold the chromatin, enabling easier transcription of the modified gene sequences. This deposition of H3K4 methylation is mainly accomplished by MLL family histone methyltransferases (HMTs), particularly MLL1(Wang, Lin et al., 2009, Wang, Zhu et al., 2012). In this study, CMI was found to induce the interaction between MLL1 and NLRP3 in microglial nuclei. Interestingly, the interaction did not lead to the activation of the NLRP3 inflammasome: IL-1β production remained unchanged in microglia after a single CMI. Instead, the interaction was related to an increase in expression of H3K4me1 and H3K4me3. Furthermore, a knockout of microglia NLRP3 significantly inhibited MLL1 elevation, suggesting that NLRP3 mediated CMI-induced trained immunity through the interaction with MLL1, independently of inflammasomes. Indeed, NLRP3 has been reported to have several inflammasome-independent functions in immunity, although IL-1β and IL-18 secretion from activated NLRP3 inflammasomes are important for various innate and adaptive immune responses. For example, NLRP3 can interact with both DNA and the transcription factor IRF4 to mediate the differentiation of Th2 cells(Bruchard, Rebe et al., 2015).

NLRP3 has been considered as a therapeutic target in the treatment of cerebrovascular diseases(Hong et al., 2019, Yang, Wang et al., 2014, Yang-Wei Fann, Lee et al., 2013). However, it is debated whether NLRP3 inhibition can reduce infarct volume in experimental ischemic strokes(Lemarchand, Barrington et al., 2019). Previously, we found that NLRP3 exacerbated the risk of recurrent ischemic strokes following a single infarction in an inflammasome-dependent manner and the data from this study further supports this hypothesis. Therefore, NLPR3 is a potential therapeutic candidate to manage the risk of recurrent ischemic strokes, depending on the degree, or severity of the initial stroke.

Nevertheless, there are some limitations to our study. Further research is required to address the function of caspase-1 and ASC in microglial nuclei. Furthermore, researchers should also investigate the other functional roles of NLRP3 in the nuclei of inflammatory cells. Moreover, microinfarcts are just one type of vascular diseases and the influence of other SVD on recurrent stroke risk requires further research.

Our study is the first to reveal the epigenetic regulation role of NLRP3 in innate immune memory, as it increases H3K4me1 and H3K4me3 through its interaction with the MLL1 complex. This innate immune memory formation causes a higher risk of recurrent strokes in mice with CMI. Therefore, NLRP3 may provide an alternative therapeutic target to mitigate the detrimental effect of CMI and subsequent stroke risk.

## Materials and Methods

### Animals

C57BL/6J male mice (6–8 weeks old, 20–25 g) were provided by the Sun Yat-sen University Medical Experimental Animal Center (Guangzhou, China). Animals were kept in a temperature and humidity-controlled specific-pathogen-free laboratory with a 12/12 h light/dark cycle. Animal experimental procedures were performed following the guidelines set by the Sun Yat-sen University Committee on the Care and Use of Animals.

### Microinfarct model

The microinfarct model was performed as previously described(Yang-Wei Fann et al., 2013). Briefly, 0.1ml of 0.3% 2000 KD FITC-dextran (FD20, Sigma) was injected into the tail vein. Under 25x magnification water-immersion, a PA measuring 15-20um in diameter and 10 um/ms in velocity, was selected as our target vessel for occlusion. Bleach mode with maximum laser power was used to damage the endothelium and bleach points were located within the vessel lumen. Irradiation was started with the 800 nm laser and was continued until a clot was visualized and the motion of RBCs was stalled. An hour’s observation was required to observe the redistribution and stabilization of the downstream flow. Irradiation was repeated if any recanalization occurred, and models with hemorrhages or diffuse burns were discarded.

### Photothrombotic stroke model

After being anesthetized with a combination of ketamine (0.12 mg/g, i.p.) and xylazine (0.01 mg/g, i.p.), mice’s skulls were exposed by making a 2-mm incision 4 mm posterior and 3 mm left of the bregma is left for further investigation. Rose Bengal (10 mg/ml, dissolved in saline) was injected in the tail vein at a dose of 0.03 mg/g body weight. The skull was then illuminated for 10 mins by two-photon with blue Hg light (450–500 nm) focusing on the aperture incision. After the illumination, the skin was sutured, and mice were kept on a warm heating pad to recover.

### Microglia isolation

The isolation of microglia was performed as previously described(Sarkar, Malovic et al., 2017). Briefly, mice were anesthetized, and brains were sampled in ice-cold PBS as quickly as possible. Brains were carefully minced into small fragments and digested by trypsin at 37 °C for 20 mins. The digested tissue was centrifuged at 400g for 5 min, centrifuged cells are then collected and resuspended in 37% percoll, add the resuspended tissue into 30% percoll and then add 70% percoll on the above layer. The tube was then centrifuged once more (700 g) for 10 min continuously. The cells between the 30 and 37% percoll were collected and resuspended in DPBS (~2×10^7^/ml). Cd11b^+^ positive magnetic isolation and was then performed with a STEMCELL Easysep isolation kit. Isolated cells were lysed in a RIPA buffer and subjected to western-blot analysis.

### Nuclear and cytoplasmic protein extraction

Isolated Cd11b^+^ microglia were further processed to extract nuclear and cytoplasmic proteins. The isolation protocol was performed according to the manufacture’s instruction (Thermoscientific). Briefly, cells were resuspended in 50 μl of lysis buffer (Thermoscientific) and incubated for 30 min on ice. Digested cells were lysed via ten strokes through a 26-gage needle and centrifuged for 10 min at 1000 g. The supernatant contained the cytoplasm and mitochondria, while the pellet contained the nucleus.

### ChIP-qPCR

Chromatin precipitation was performed as previously described(Bertani, Kan et al., 2008). Briefly, isolated microglia were fixed in 0.75% formaldehyde for 10 min and quenched using glycine. Chromatin was then sonicated for four cycles of 10X (30 s ON, 30 s OFF) at 10% maximum power. Chromatin precipitation was performed using a rabbit anti-mouse H3K4me3 IgG Ab (Cell Signaling Technology). DNA was then quantified using qPCR with the following primer pairs:

GAPDH, FW 59-ATCCAAGCGTGTAAGGGTCC-39, RV 59-GACTGAGATTGGCCCGATGG-39; IL-6, FW 59-AGCTCTATCTCCCCTCCAGG-39, RV 59-ACAC-CCCTCCCTCACACAG-39; TNF-α, FW 59-CAGGCAGGTTCTCTTCCTCT-39, RV 59-GCTTTCAGTGCTCATGGTGT-39.

For all ChIP experiments, qPCR values were normalized as the percentage recovery of the input DNA.

### Western-Blot analysis

Mice were euthanized and perfused with ice-cold saline. The ischemic cortex was sampled and homogenized by ultrasonic in RIPPA. Equal mass of protein quantified by BCA assay was subjected to western-blot analysis.

The following primary antibodies were used: anti-β-actin (1:1000; cell-signaling technology, USA), anti-IL-1β (1:1000; cell-signaling technology, USA), anti-IL-6 (1:1000; cell-signaling technology, USA), anti-H3K4me1 (1:1000; cell-signaling technology, USA), anti-H3K4me1 (1:1000; cell-signaling technology, USA), and anti-H3 (1:1000; cell-signaling technology, USA). The following secondary antibodies used were goat-anti-rabbit HRP-linked antibody (1:1000; cell-signaling technology, USA), goat-anti-mouse HRP-linked antibody (1:1000; cell-signaling technology, USA).

### Immunofluorescence and in situ proximity ligation assay

Mice were euthanized, then perfused with ice-cold saline and fixed using 4% formaldehyde. After dehydration by 30% sucrose, coronal brain slices (20-μm thick) from the right parietal cortices were sectioned using a frozen microtome (Leica) at intervals of 200 μm to produce consecutive frozen sections.

For immunofluorescence experiments, slices were first permeabilized by 0.1% Triton X-100 in PBS for 15 min, then blocked with PBS containing 5% BSA for one hour. Primary antibodies diluted in blocking buffer were added to the slices and incubated at 4 °C overnight. Slices were washed three times with PBS and incubated for 30 min at room temperature with the secondary antibody, then washed three more times with PBS. Slices were finally mounted with a DAPI mounting medium (Sigma-Aldrich) for subsequent observations.

For proximity ligation assay experiments, after primary antibodies were washed out, cells were incubated for 1 h at 37 °C with the appropriate probes and were washed twice with PBS. Probes were then ligated for 30 min at 37 °C and washed twice in buffer A, then amplified for 100 min at 37 °C in dark conditions with polymerase (Sigma-Aldrich).

The following primary antibodies were used in this experiment: anti-iba1 (1:500; abcam), anti-NLRP3 (1:200; Thermoscientific), anti-MLL1 (1:100; cell-signaling technology, USA) The following secondary antibodies were used: goat-anti-rabbit HRP-linked antibody (1:1000; cell-signaling technology, USA), goat-anti-mouse HRP-linked antibody (1:1000; cell-signaling technology, USA).

### Nissls staining

Nissls staining was performed using serial frozen coronal sections, 10 μm in thickness and 200 μm in interval from PPC were used. The sections were hydrated in 1% toluidine blue at 37 °C overnight and washed out by double distilled water. After soaking in dimethylbenzene for 5 s, sections were sealed in permount covered by coverslip. Infarct volume (mm^3^) was used to measure the infarct size by the Image J.

### ELISA

Blood serum was obtained by coagulating mice blood in vacuettes (Bio-rad) for 10 min at room temperature, then centrifuging it for 10 min at 2,000 g. Serum samples were diluted 1:2 before measurements. Blood serum was then assayed using ELISA for IL-6, IL-1β, and TNF-α (R&D Systems) according to the manufacturers’ instructions.

### Statistical analysis

Data were analyzed with Graphpad Prism 7.0. and represented as mean ± standard deviation (SD). Different treatment groups were evaluated using t-test or two-way ANOVA with Turkey’s test for multiple comparisons to determine differences between individual groups. The significance level was set at *p* < 0.05. Regardless of the method used, the results are equivalent in magnitude and statistically significant. *For priori* sample size was calculated based on power as 0.95 and α as 0.05.

## Acknowledgements

This study was supported by grants from the National Natural Science Foundation of China (No. 81873751, No.81671102), the National Key Research and Development Program of China, Stem Cell and Translational Research (No. 2017YFA0105104), Guangdong provincial science and technology plan project (No. 2016B030230002), the Southern China International Cooperation Base for Early Intervention and Functional Rehabilitation of Neurological Diseases (2015B050501003), Guangdong Provincial Engineering Center For Major Neurological Disease Treatment, Guangdong Provincial Translational Medicine Innovation Platform for Diagnosis and Treatment of Major Neurological Disease, Guangdong Provincial Clinical Research Center for Neurological Diseases.

## Author Contributions

YW and ZP drafted the manuscript. TW, YK and YG accomplished the experiment. FY and GL designed the experiment. YC, YL and SH provided technical support. ZY, GQ, ZP provided financial support.

---

**Fig. S1.**
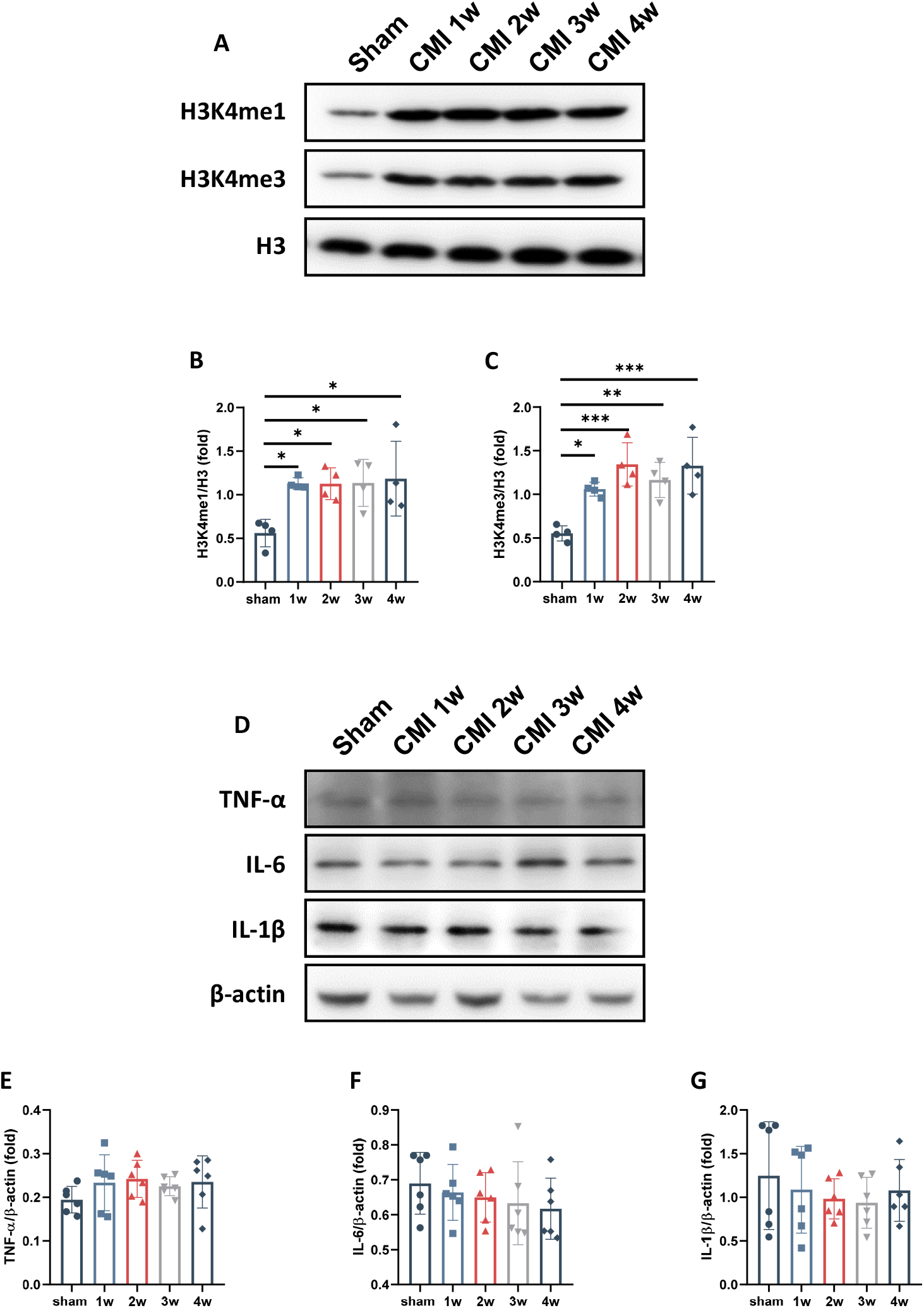
Quantitative analysis of western-blot data in Fig1. A-C Immunoblots (A) and quantitative analysis of related H3K4me1 and H3K4me3 expression (B, C) D-G Immunoblots (D) and quantitative analysis of TNF-α, IL-6, and IL-1β expression (E-G).

**Fig. S2.**
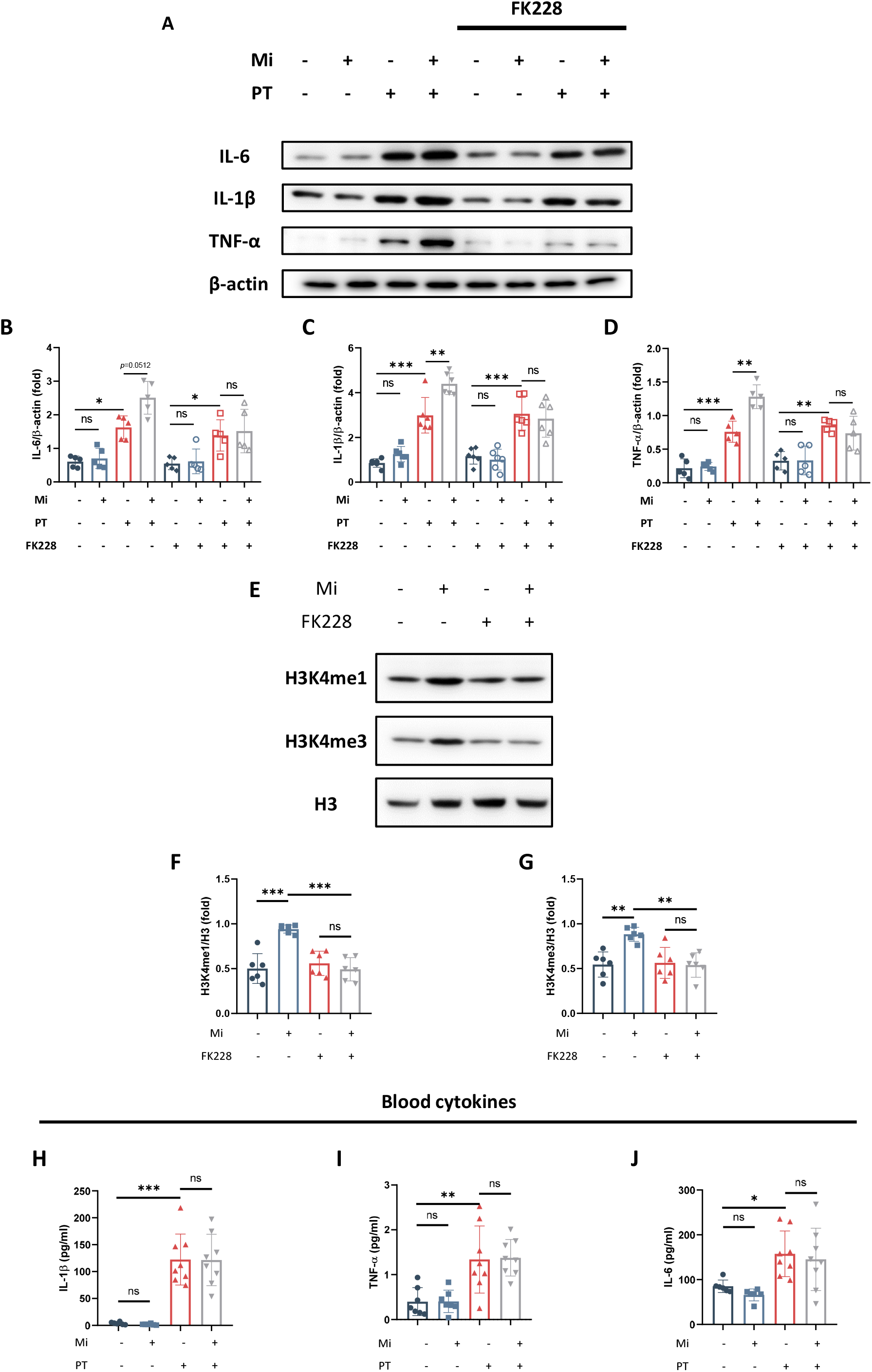
Quantitative analysis of western-blot data in Fig2. A-D Immunoblots (A) and quantitative analysis of TNF-α, IL-6, and IL-1β expression (B-D). E-G Immunoblots (E) and quantitative analysis of related H3K4me1 and H3K4me3 expression (F, G). H-J ELISA showed that stroke mice with preceding microinfarct did not deteriorate the pro-inflammatory cytokines in the peripheral blood.

**Fig. S3.**
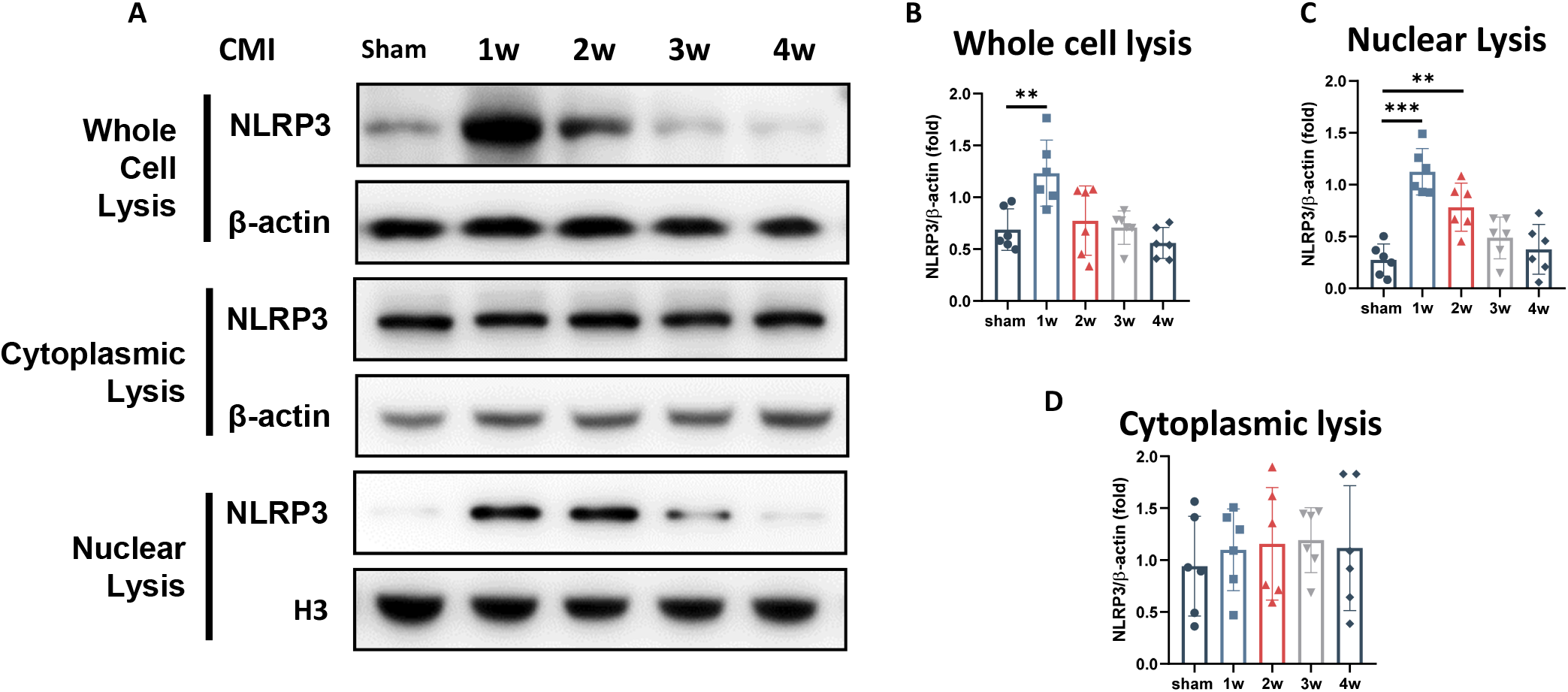
Quantitative analysis of western-blot data in Fig 3. A-D Immunoblots (A) and quantitative analysis of NLRP3 expression in the whold-cell lysis (B), Nuclear lysis (C) and Cytoplasmic lysis (D)

**Fig. S4.**
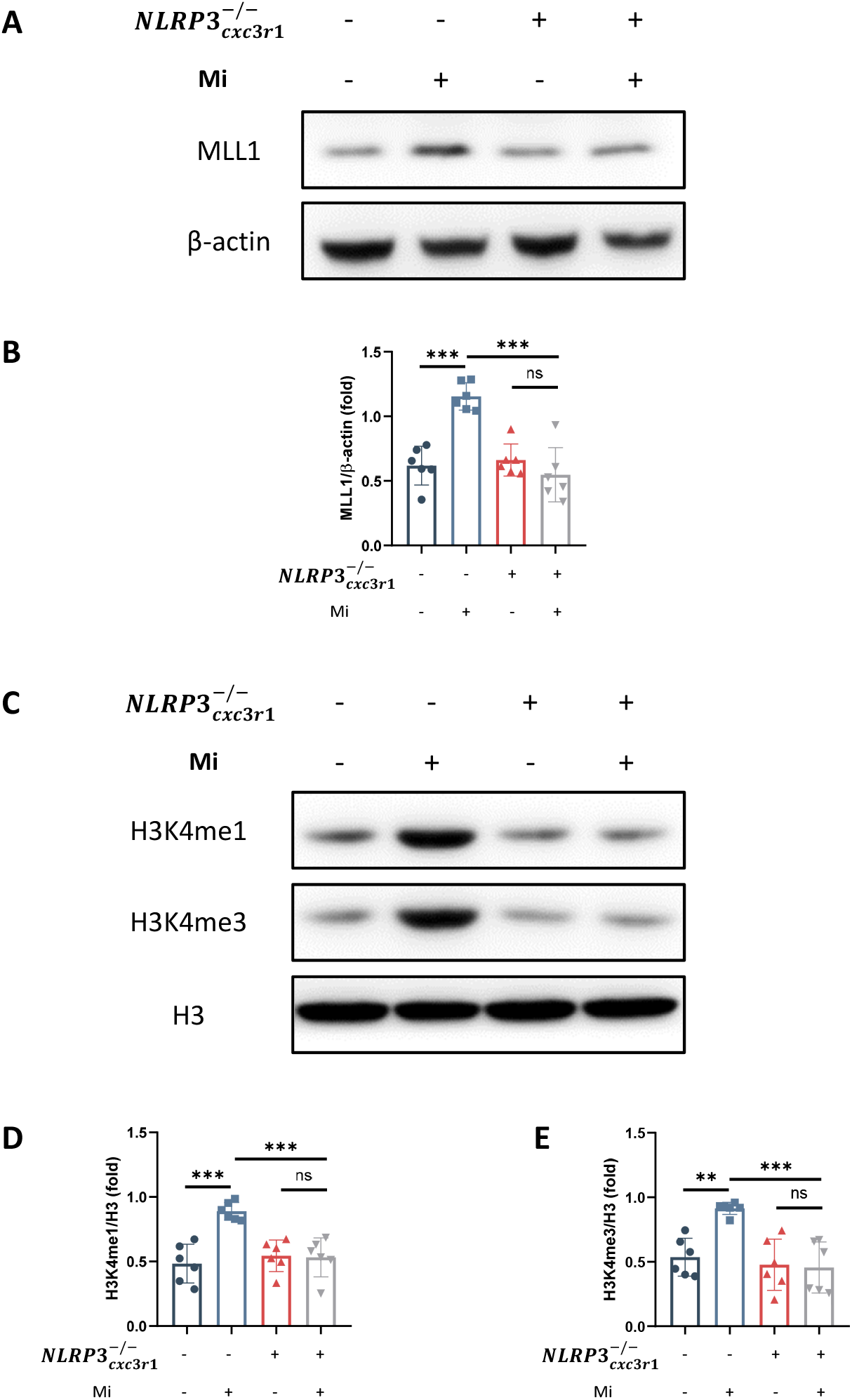
Quantitative analysis of western-blot data in Fig 4. A-B Immunoblots (A) and quantitative analysis of MLL1 expression (B). C-E Immunoblots (C) and quantitative analysis of H3K4me1 (D) and H3K4me3 expression (E).

